# LSMMD-MA: Scaling multimodal data integration for single-cell genomics data analysis

**DOI:** 10.1101/2022.03.23.485536

**Authors:** Laetitia Meng-Papaxanthos, Ran Zhang, Gang Li, Marco Cuturi, William Stafford Noble, Jean-Philippe Vert

## Abstract

**Motivation:** Modality matching in single-cell omics data analysis—i.e., matching cells across data sets collected using different types of genomic assays—has become an important problem, because unifying perspectives across different technologies holds the promise of yielding biological and clinical discoveries. However, single-cell dataset sizes can now reach hundreds of thousands to millions of cells, which remains out of reach for most multi-modal computational methods.

**Results:** We propose LSMMD-MA, a large-scale Python implementation of the MMD-MA method for multimodal data integration. In LSMMD-MA we reformulate the MMD-MA optimization problem using linear algebra and solve it with KeOps, a CUDA framework for symbolic matrix computation in Python. We show that LSMMD-MA scales to a million cells in each modality, two orders of magnitude greater than existing implementations.

**Availability:** LSMMD-MA is freely available at https://github.com/google-research/large_scale_mmdma

## 1 Introduction and background

Modality matching in single-cell genomics data analysis can enhance our understanding of the relationships between cellular modalities and help us resolve cell states. In this problem, single-cell measurements collected using two or more different types of assays are projected into a shared space or are otherwise matched across modalities, with the goal of achieving insights into the joint multi-modal dataset. Most existing multimodal models rely on learning cell representations in each modality in a joint low-dimensional space [1, 2, 3, 4, 5, 6, 7, 8]. MMD-MA [9, 10] is one such method that has shown promising results on datasets containing several thousand cells in each modality. However, thanks to new single-cell technologies, the size of single-cell datasets has increased significantly in the past two years, now reaching several hundreds of thousands to millions of cells [11, 12, 6]. These datasets cannot be analyzed by current implementations of MMD-MA due to memory issues.

More precisely, MMD-MA [9] is a multimodal approach that maps each cell in each modality to a shared, low-dimensional representation space. The linear mappings from the input spaces to the representation space are learned by minimizing an objective function composed of several terms: a) a *matching* term based on the squared maximum mean discrepancy (MMD) with a Gaussian radial basis function (RBF) kernel to ensure that the different modalities overlap in the representation space, b) two *non-collapsing* penalties to prevent trivial solutions, and c) two *distortion* penalties to ensure that as much information from the input data as possible is captured in the shared representation. More details about the method are provided Appendix A. However, current implementations of MMD-MA [9, 10] scale quadratically as a function of the number of cells in memory and runtime, which is prohibitive for datasets with more than a few thousand samples (see Table A1).

To increase the scalibility of MMD-MA, we introduce LSMMD-MA, a reformulation and PyTorch implementation of MMD-MA that overcomes the memory explosion issue. To achieve this, we 1) reformulate MMD-MA’s optimization problem in the primal, which is beneficial when the number of cells is larger than the number of informative features, and 2) implement the MMD matching term with the CUDA-based KeOps library for symbolic matrices [13], tailored to handle matrices that do not fit in RAM or GPU memory. The resulting algorithm scales only linearly in memory with the number of cells and can handle up to a million cells in each modality.

## 2 Methods

### Reformulating MMD-MA in the primal

Let 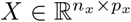 (resp. 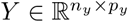) be the data matrix of the first (resp. second) modality, where *n*_*x*_ (resp. *n*_*y*_) is the number of cells in the first (resp. second) modality and *p*_*x*_ (resp. *p*_*y*_) is the number of features in the first (resp. second) modality. The goal of MMD-MA is to learn two mappings from the input spaces 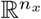 and 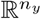 to a shared representation space ℝ^*d*^. We focus specifically on linear mappings, as in the original publications [9, 10], where mappings are parameterized with dual variables 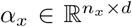 and 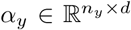 such that the embedding of the first (resp. second) modality is *XX*^⊤^*α*_*x*_ (resp. *YY*^⊤^*α*_*y*_). Instead, we equivalently parameterize the mappings by primal variables 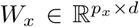 and 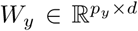, such that the embedding of the first (resp. second) modality is *XW*_*x*_ (resp. *YW*_*y*_). We can then rewrite the MMD-MA optimization problem in the primal:

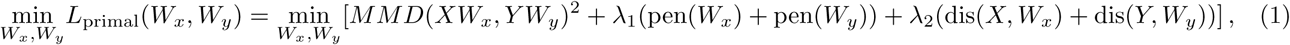

where *λ*_1_ and *λ*_2_ are hyperparameters. Under the assumption that *n >> p >> d* for each modality, efficiently implementing the primal loss (1) scales better than implementing the dual loss, as shown in Table A1 shows. The primal loss does not require the computation and storage of the linear kernel matrices *XX*^⊤^ and *YY* ^⊤^, which are O(*n*^2^) in time and memory, and the penalty and distortion terms are not O(*n*^2^) in runtime anymore. However, computing the MMD term remains O(*n*^2^) in runtime and memory if we implement it naively.

### Using KeOps

To overcome the O(*n*^2^) memory burden of computing the MMD term, we implement it using the CUDA-based Map-Reduce scheme of KeOps [13]. This allows us to compute the MMD term without instantiating the *n* × *n* Gaussian RBF kernel in memory, using symbolic matrix computation with O(*n*) memory complexity. KeOps therefore optimizes (1) with a linear memory complexity and also improves runtime by a significant multiplicative factor when the number of samples is larger than 1000.

## 3 Implementation

We make four algorithms available, including `LS-MMDMA` and three variants: primal formulation without KeOps, dual formulation with KeOps, and dual formulation without KeOps (an efficient implementation of the original algorithm). The code is implemented in PyTorch [14] and can run on CPU or GPU. The package is open source with an Apache license, available at github.com/google-research/large_scale_mmdma. It is referenced on PyPI and can be installed with the command: `pip install lsmmdma`. Details about I/O, command line instructions and tutorials are given in the `Readme.md` and in the `examples` folder.

## 4 Results and conclusion

We tested the implementation of LSMMD-MA (primal formulation with KeOps) against three comparison partners (primal formulation without KeOps, dual formulations with KeOps and dual formulation without KeOps) and against the two original implementations [9, 10] which focus on the dual formulation in TensorFlow [15] and PyTorch [14], respectively. Additionally, [10] proposes to use the linear time approximation of MMD for large numbers of samples (*>* 5000) (see Lemma 14 in [16]). We ran all algorithms on simulated datasets of different sizes where the latent space is shaped as a branch (see lsmmdma/data/data_pipeline.py in GitHub and the Appendix D for more details). All algorithms were run for 500 epochs and a low-dimensional representation of dimension *d* = 10. A V100 GPU (16GB) was used for the experiments.

We observe that LSMMD-MA, using the primal formulation and KeOps, scales to one million cells in each modality, whereas the original implementations runs out of memory for more than 14,000 cells (Figure 1). We also notice that our dual implementations are faster than the original implementations irrespective of the number of samples. As a proof-of-principle, we also ran LSMMD-MA on a real-world CITE-seq dataset containing 90,261 human bone marrow mononuclear cells, with 13,953 gene IDs for the gene expression modality and 134 proteins for the protein marker modality [17]. This dataset required 68.4 minutes to run 1000 epochs, which would have been infeasible with previous versions of MMD-MA.

**Figure 1:**
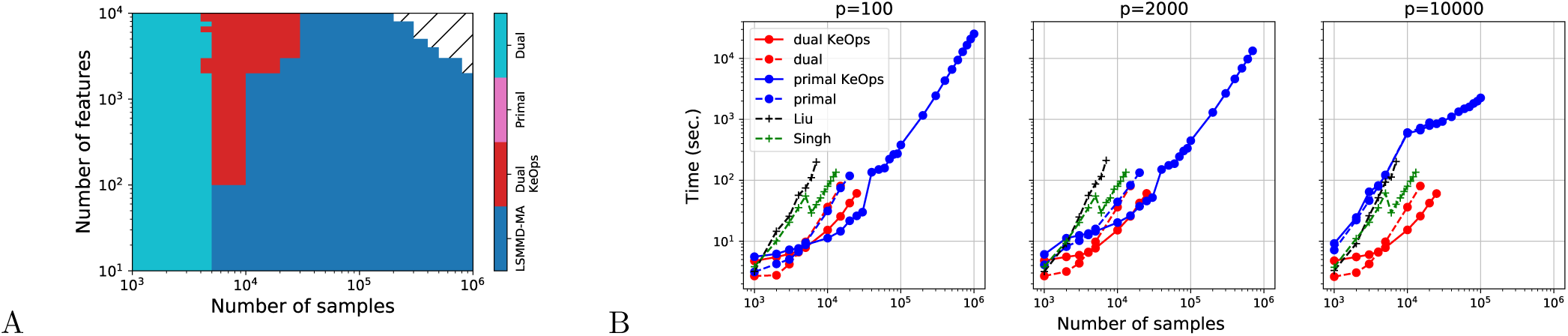
(A) Fastest MMD-MA variant as a function of the number of samples and the number of features. Hatched region means that all variants ran out of memory. (B) Runtime as a function of number of cells for different implementations of MMD-MA, when the dimension *p* of the input data varies. The runtime for a larger set of *p* is shown Figure A1.

These results suggest that an optimised implementation, exploiting the primal formulation and taking advantage of the KeOps library, are key to building a multimodal model that scales to the size of current single-cell datasets.

## Appendix

### A Description of MMD-MA with the dual formulation

`MMD-MA` [9] is a method for analyzing multimodal data that relies on mapping the observed cell samples to embeddings, using functions belonging to a reproducing kernel Hilbert space (RKHS). The authors build linear kernels for both domains and learn the coefficients of the embedding functions in the RKHS dual representation. In practice, the `MMD-MA` loss is composed of a) the squared MMD, a *matching* term, b) two *non-collapsing* penalties and c) two *distortion* penalties, one for each modality.

Let 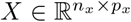 be the data matrix of the first modality and 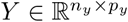 the one of the second modality. *n*_*x*_ (resp. *n*_*y*_) is the number of cells in the first (resp. second) modality and *p*_*x*_ (resp. *p*_*y*_) is the number of features in the first (resp. second) modality. We denote by 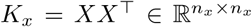 and 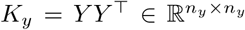 the linear kernel matrices corresponding to both datasets. 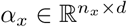 and 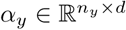 are the learned coefficients of the RKHS functions used to build the embeddings *K*_*x*_*α*_*x*_ and *K*_*y*_*α*_*y*_, of dimension *d*. The loss of `MMD-MA` can be written as follows:

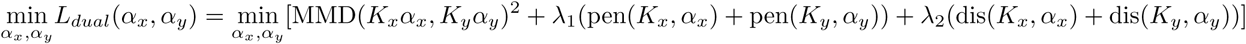

where *λ*_1_ and *λ*_2_ are two hyperparameters weighing the non-collapsing penalty (*pen*(*K, α*) = ||*α*^*T*^ *Kα* − **I**_*d*_ ||_2_) and the distortion penalty, ensuring that the geometries in the low- and high-dimensional representations are comparable (*dis*(*K, α*) = ||*K* −*K*^*T*^ *α*^*T*^ *αK* ||_2_). The RBF kernel is used to calculate MMD, introducing non-linearity in the matching term. However, the linear and gaussian kernel matrices require memory and runtime that scale quadratically as a function of the number of cells in terms of memory and runtime, which is prohibitive for large datasets.

### B Runtime and memory requirements

**Table A1:** Runtime and memory requirements for MMD-MA in the primal and dual forms without using KeOps. *n* is the number of cells, *p* the number of features and *d* the dimension of the embeddings. We remove the subscripts *x* and *y* for readability but the O notations of the table hold for each modality. Typically, *n >> p >> d* for each modality.

| Model | MMD-MA dual |  |  |  | MMD-MA primal |  |  |  |
| --- | --- | --- | --- | --- | --- | --- | --- | --- |
|  | input | MMD | pen | dis | input | MMD | pen | dis |
| Runtime | $O(\mathbf{n}^2 p)$ | $O(\mathbf{n}^2 d)$ | $O(\mathbf{n}^2 d + nd^2)$ | $O(\mathbf{n}^2 d)$ | - | $O(\mathbf{n}^2 d + pnd)$ | $O(d^2 p + d^2)$ | if $n > p$: $O(p^2 n + p^3 + p^2 d)$<br>else: $O(n^2 p + nd)$ |
| Memory | $O(\mathbf{n}^2)$ | $O(\mathbf{n}^2)$ | $O(nd + d^2)$ | $O(nd)$ | $O(np + pd)$ | $O(\mathbf{n}^2 + nd)$ | $O(d^2)$ | $O(\min(p, n) * (p + n + d))$ |

### C Benchmark

**Figure A1:**
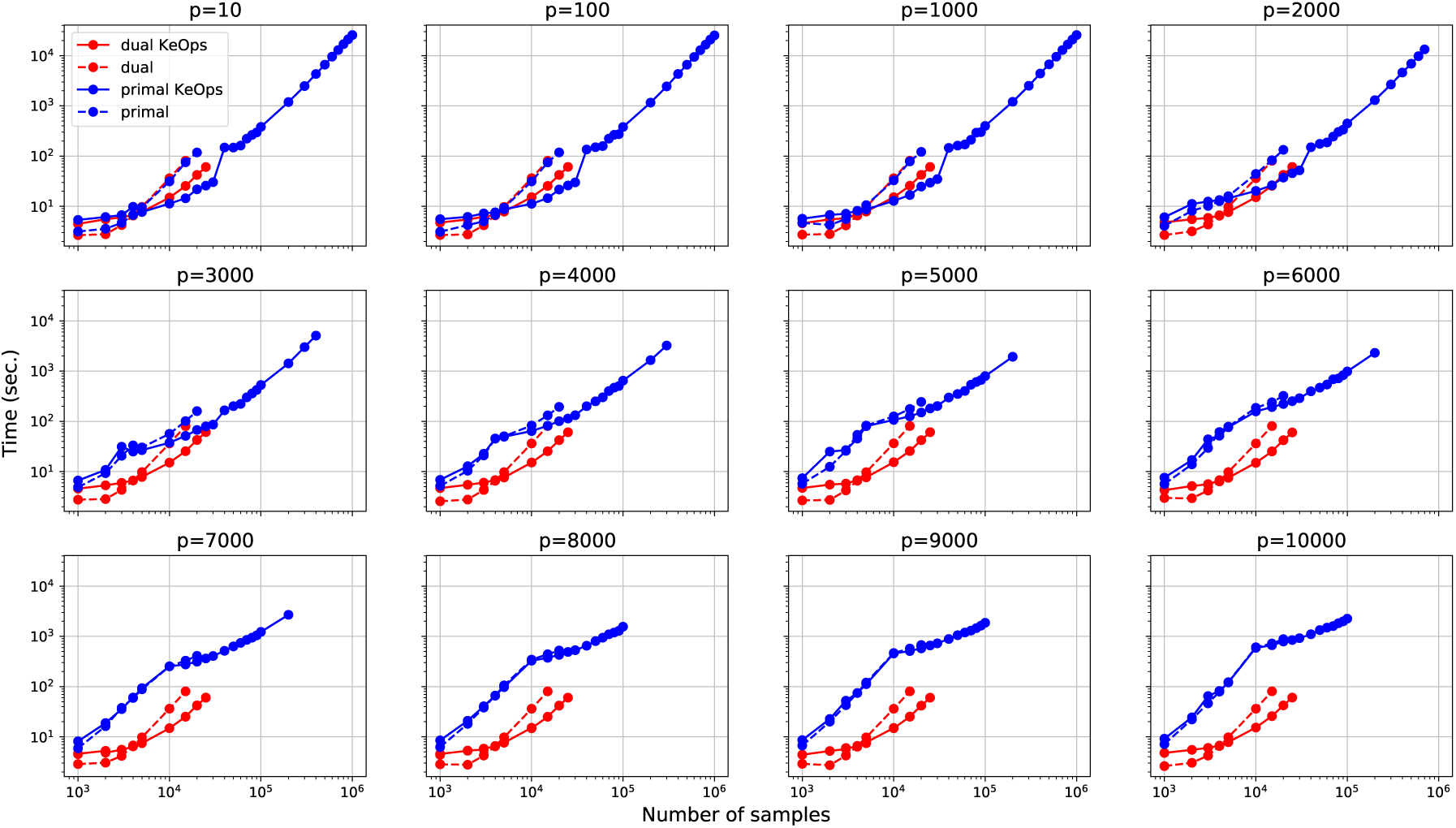
Runtime as a function of number of cells for different implementations of MMD-MA, when the dimension *p* of the input data varies.

### D Datasets

The synthetic datasets used for the runtime experiments have a varying number of samples from 10^3^ to 10^6^ and a varying number of features from 10 to 10^4^. To generate them, we used the `generate_data` function in lsmmdma/data/data_pipeline.py, with the `random_seed` argument fixed to 4 and the `simulation` argument set to ‘branch’. The simulation process works as follows. A manifold (of shape ‘branch’) is generated in two dimensions. The resulting set of points is standardised. The points are then mapped to `p_feature` dimensions using a (2 x `p_feature`) mapping, sampled from a standard Gaussian distribution, resulting in a (`n_sample` x `p_feature`) matrix. Gaussian noise is then added to each element of the matrix. This simulation was first described in [9] and the latent space shaped as a branch aims at mimicking a development process for example.

The real-world dataset on which the last experiment is ran comes from [17], where datasets were made publically available for the [Neurips Competition](https://openproblems.bio/neurips_2021/) Multimodal Single-Cell Data Integration.

## References

[1] Cao, K., Bai, X., Hong, Y., and Wan, L. (2020). Unsupervised topological alignment for single-cell multi-omics integration. Bioinformatics 36, i48–i56. doi:10.1093/bioinformatics/btaa443.

[2] Gayoso, A., Steier, Z., Lopez, R., Regier, J., Nazor, K.L., Streets, A., and Yosef, N. (2021). Joint probabilistic modeling of single-cell multi-omic data with totalVI. Nat. Methods 18, 272–282. doi:10.1038/s41592-020-01050-x.

[3] Jin, S., Zhang, L., and Nie, Q. (2020). scAI: an unsupervised approach for the integrative analysis of parallel single-cell transcriptomic and epigenomic profiles. Genome Biol. 21, 1–19. doi:10.1186/s13059-020-1932-8.

[4] Stark, S.G., Ficek, J., Locatello, F., Bonilla, X., Chevrier, S., Singer, F., Consortium, T.P., Rätsch, G., and Lehmann, K.V. (2020). SCIM: Universal single-cell matching with unpaired feature sets. Bioinformatics 36, i919– i927. doi:10.1093/bioinformatics/btaa843.

[5] Welch, J.D., Kozareva, V., Ferreira, A., Vanderburg, C., Martin, C., and Macosko, E.Z. (2019). Single-cell multi-omic integration compares and contrasts features of brain cell identity. Cell 177, 1873–1887. doi:10.1016/j.cell.2019.05.006.

[6] Hao, Y., Hao, S., Andersen-Nissen, E., Mauck III, W.M., Zheng, S., Butler, A., Lee, M.J., Wilk, A.J., Darby, C., Zager, M., et al. (2021). Integrated analysis of multimodal single-cell data. Cell 184, 3573–3587.

[7] Cao, Z.J. and Gao, G. (2021). Multi-omics integration and regulatory inference for unpaired single-cell data with a graph-linked unified embedding framework. bioRxiv.

[8] Raimundo, F., Meng-Papaxanthos, L., Vallot, C., and Vert, J.P. (2021). Machine learning for single-cell genomics data analysis. Current Opinion in Systems Biology 26, 64–71.

[9] Liu, J., Huang, Y., Singh, R., Vert, J.P., and Noble, W.S. (2019). Jointly embedding multiple single-cell omics measurements. In 19th International Workshop on Algorithms in Bioinformatics (WABI 2019), Leibniz International Proceedings in Informatics (LIPIcs), volume 143. pp. 10:1–10:13. doi:10.4230/LIPIcs.WABI.2019.10.

[10] Singh, R., Demetci, P., Bonora, G., Ramani, V., Lee, C., Fang, H., Duan, Z., Deng, X., Shendure, J., Disteche, C., et al. (2020). Unsupervised manifold alignment for single-cell multi-omics data. In Proceedings of the 11th ACM International Conference on Bioinformatics, Computational Biology and Health Informatics. pp. 1–10. doi:10.1101/2020.06.13.149195.

[11] Rozenblatt-Rosen, O., Shin, J.W., Rood, J.E., Hupalowska, A., Regev, A., and Heyn, H. (2021). Building a high-quality human cell atlas. Nature Biotechnology 39, 149–153.

[12] Papatheodorou, I., Moreno, P., Manning, J., Fuentes, A.M.P., George, N., Fexova, S., Fonseca, N.A., Füllgrabe, A., Green, M., Huang, N., et al. (2020). Expression atlas update: from tissues to single cells. Nucleic acids research 48, D77–D83.

[13] Charlier, B., Feydy, J., Glaunés, J.A., Collin, F.D., and Durif, G. (2021). Kernel operations on the gpu, with autodiff, without memory overflows. Journal of Machine Learning Research 22, 1–6.

[14] Paszke, A., Gross, S., Massa, F., Lerer, A., Bradbury, J., Chanan, G., Killeen, T., Lin, Z., Gimelshein, N., Antiga, L., et al. (2019). Pytorch: An imperative style, high-performance deep learning library. In Advances in Neural Information Processing Systems 32, H. Wallach, H. Larochelle, A. Beygelzimer, F. d’Alché-Buc, E. Fox, and R. Garnett, eds. (Curran Associates, Inc.), pp. 8024–8035.

[15] Abadi, M., Agarwal, A., Barham, P., Brevdo, E., Chen, Z., Citro, C., Corrado, G.S., Davis, A., Dean, J., Devin, M., et al. (2015). TensorFlow: Large-scale machine learning on heterogeneous systems. Software available from tensorflow.org.

[16] Gretton, A., Borgwardt, K.M., Rasch, M.J., Schölkopf, B., and Smola, A. (2012). A kernel two-sample test. The Journal of Machine Learning Research 13, 723–773.

[17] Luecken, M.D., Burkhardt, D.B., Cannoodt, R., Lance, C., Agrawal, A., Aliee, H., Chen, A.T., Deconinck, L., Detweiler, A.M., Granados, A.A., et al. (2021). A sandbox for prediction and integration of dna, rna, and proteins in single cells. In Thirty-fifth Conference on Neural Information Processing Systems Datasets and Benchmarks Track (Round 2).

